# Remodeling of the *C. elegans* Non-coding RNA Transcriptome by Heat Shock

**DOI:** 10.1101/706820

**Authors:** William P. Schreiner, Delaney C. Pagliuso, Jacob M. Garrigues, Jerry S. Chen, Antti P. Aalto, Amy E. Pasquinelli

**Affiliations:** Division of Biology, University of California, San Diego, La Jolla, CA 92093-0349, USA; Encoded Genomics, South San Francisco, CA 94080, USA; ProQR Therapeutics N.V., 2333 CK Leiden, The Netherlands

**Keywords:** heat shock, HSF-1, *C. elegans*, miRNA, lincRNA, Helitron

## Abstract

Elevated temperatures activate a Heat Shock Response (HSR) to protect cells from the pathological effects of protein mis-folding, cellular mis-organization, organelle dysfunction and altered membrane fluidity. This response includes activation of the conserved transcription factor Heat Shock Factor 1 (HSF-1), which binds Heat Shock Elements (HSEs) in the promoters of genes induced by heat shock (HS). The up-regulation of protein-coding genes (PCGs), such as Heat Shock Proteins (HSPs) and cytoskeletal regulators, is critical for cellular survival during elevated temperatures. While the transcriptional response of PCGs to heat shock has been comprehensively analyzed in a variety of organisms, the effect of this stress on the expression of non-coding RNAs (ncRNAs) has not been systematically examined. Here we show that in *Caenorhabditis elegans* HS induces up- and down-regulation of specific ncRNAs from multiple classes, including miRNA, piRNA, lincRNA, pseudogene, and repeat elements. Moreover, some ncRNA genes appear to be direct targets of the HSR, as they contain HSF-1 bound HSEs in their promoters and their expression is regulated by this factor during HS. These results demonstrate that multiple ncRNA genes respond to HS, some as direct HSF-1 targets, providing new candidates that may contribute to organismal survival during this stress.

## INTRODUCTION

In the natural world, animals experience a variety of potentially lethal environmental challenges. Heat stress is one of the most recognizable and can be fatal, if not properly addressed. Organisms have evolved an ancient response to cope with this dangerous insult: The Heat Shock Response (HSR) (1). Heat Shock Factor 1 (HSF-1), serves as a master transcriptional regulator of the HSR in eukaryotes (2). Elevated temperatures trigger activation and binding of HSF-1 to Heat Shock Elements (HSEs) in the promoters of genes encoding Heat Shock Proteins (HSPs) and other factors that protect the cell from heat-induced damage. For example, HSPs act as molecular chaperones to deal with the rampant protein misfolding associated with heat stress. While heat shock has been shown to induce widespread changes in gene expression in cell as well as animal models, a limited set of genes appears to be direct targets of HSF-1 activation (3–5). Instead, downstream transcriptional effectors mediate many of the other HS-induced changes in gene expression (3–5).

There is also potential for post-transcriptional mechanisms to regulate the levels of protein-coding mRNAs during the HSR (6–9). In *C. elegans*, specific microRNAs (miRNAs) have been found to be up- or down-regulated during HS (10, 11). MiRNAs function as small, ~22 nucleotide, guide RNAs that use partial base-pairing to recruit Argonaute (AGO) proteins to specific mRNAs, triggering mRNA destabilization and translational repression (12). While direct mRNA targets of HS-regulated miRNAs are yet to be determined, loss of some of these miRNAs has been shown to affect the viability of *C. elegans* subjected to HS (11, 13). For example, deletion of *miR-71* or *miR-239* results in reduced or enhanced survival at elevated temperatures, respectively.

In addition to miRNAs, the expression of other types of non-coding RNAs (ncRNAs) can respond to fluctuations in environmental conditions (14, 15). The *C. elegans* long ncRNA, *rncs-1* (RNA non-coding starvation up-regulated) is induced by food deprivation and potentially regulates other small RNA pathways (15). In a variety of organisms, HS and other stress conditions have been shown to cause accumulation of RNAs from transposon and other types of repetitive sequences (16). Currently, it is unclear if the HS-induced changes in ncRNA expression are a direct result of the HSR or the aftermath of defective transcriptional or post-transcriptional silencing mechanisms under these conditions. Additionally, whether specific ncRNAs or generally all members of a certain class respond to HS has not been systematically investigated.

To examine the organismal response of different classes of ncRNAs to HS, we surveyed the expression of miRNAs, Piwi RNAs (piRNAs), long intergenic ncRNAs (lincRNAs), ncRNAs, pseudogene- and repeat-derived RNAs in *C. elegans* under HS versus control temperature conditions. Within each class, we observed HS-induced changes for specific transcripts that included rapid and dramatic up-regulation of some ncRNAs. Similar to canonical HS-responsive protein-coding genes, we found that the promoter sequences of the miRNA miR-239, *Helitron1_CE* transposons, and the pseudogene *dct-10* contain Heat Shock Elements (HSEs) that are bound by HSF-1 in response to HS. Furthermore, we show that up-regulation of these ncRNAs in HS is regulated by HSF-1, suggesting that they are direct transcriptional targets of the HSR. Overall, our comprehensive analysis of ncRNA expression in response to HS revealed new types of molecules that are regulated by and, in turn, may contribute to organismal survival in the face of this stress.

## MATERIALS AND METHODS

### Sequencing and analysis of mRNAs and long ncRNAs

N2 wild-type worms were grown to L4 stage in a 20°C incubator under standard growth conditions (17). The experimental group was subjected to heat stress by raising the temperature to 35°C for 4 hours. Animals were then collected, snap-frozen, and total RNA was extracted using a standard Trizol RNA extraction protocol. cDNA sequencing libraries were prepared from total RNA from N2 wild-type control or heat shocked worms using the standard protocol from the Illumina Stranded TruSeq RNA library prep kit. Prior to library preparations, ribosomal RNA was removed using RiboZero Gold (Illumina). cDNA libraries were sequenced on an Illumina Genome Analyzer II (100 bp paired-end reads). FASTQ reads were first trimmed using fastq-mcf (https://expressionanalysis.github.io/ea-utils/), which removed flanking Illumina adapter sequence as well as nucleotides with low quality sequencing scores. Reads were then aligned to the *C. elegans* genome WS235 using STAR (18). Aligned reads were sorted using Samtools (19). Reads were counted using FeatureCounts and Ensembl 88 gene annotations (20). Differential expression of gene expression was determined using DESeq2 (21). Pseudogenes, lincRNAs, and ncRNAs are included in the Ensembl 88 gene annotations. After differential expression, these classes of genes were filtered out of the results and analyzed separately. See github.com/wschrein for code and additional example graphs.

To identify false positive up-regulated mRNAs that likely resulted from failure of Pol II transcriptional termination of an upstream HS-responsive gene, we used a strategy similar to that in Duarte et al., 2016 (3). First, an intron retention score (IR score) for each gene was calculated by dividing the total normalized intron reads by the total normalized exon reads per gene. Reads were normalized by DESeq2 which normalizes to sequencing depth but not length. Next, up-regulated genes were analyzed for accumulation of Intergenic Junction (IJ) reads between their annotated start site and the closest upstream gene. To do this, a 21 bp region that overlaps 11 nts into the 5’ annotated start site and 10 nts upstream of the start site was obtained for each gene. The location for these regions was obtained by parsing a list of Intergenic regions downloaded from the WS235 version of the Wormbase ftp site (22). Reads for this region in both control and heat shock samples were obtained using the program FeatureCounts. The IJ Ratio for each gene was calculated by dividing the normalized (for depth) HS Intergenic reads by the CTRL Intergenic reads. Genes with an IR score greater than 0.4 and an IJ ratio greater than 2 were removed from the list of up-regulated PCGs. In addition, PCGs with an IR score greater than 1 were also filtered out as these reads derived from independently transcribed ncRNAs, such as tRNAs and snoRNAs, present in the intron of the PCG. Finally, PCGs that overlapped a repetitive element by > 50% were filtered out. A list of *C. elegans* repetitive elements was obtained from the UCSC genome Browser. Overlap was determined using Bedtools intersect (23).

To analyze the expression of different classes of Repetitive Element RNAs, RNA-Sequencing data were aligned to a set of *C. elegans* consensus repeats obtained from repbase (24). Primary aligned reads were obtained using the following command: samtools view -F 260 ${s} | cut -f 3 | sort | uniq -c | awk ‘{printf(“%s\t%s\n”, $2, $1)}’ > ${s}counts.txt Differential expression was determined using DESeq2. More detailed information including sample code/commands can be found on github.com/wschrein.

### Small RNA sequencing and analysis

*C. elegans* were grown to the L4 stage at 20°C then shifted to 35°C for six hours. Animals were then collected, snap-frozen, and total RNA was extracted using a standard Trizol RNA extraction protocol. Small-RNA libraries were prepared using Illumina’s TruSeq Small RNA library prep kit. miRNAs were analyzed by mapping small RNA sequencing data to *C. elegans* miRNAs. DESeq2 was used to determine differential expression from two independent biological replicates. piRNAs were analyzed by mapping small RNA sequencing data to a database of *C. elegans* piRNAs obtained from Wormbase using the STAR aligner. Primary reads were obtained using samtools, and differential expression was determined using DESeq2.

For miRNA seed analysis, the longest 3’ UTR isoform for each gene was considered. UTR annotations were obtained from the WS263 GTF annotation from Wormbase. Cytoscape was used to generate the network analysis graphs (25).

### ChIP-seq data mapping, peak calling, and normalization

HSF-1 and Pol II ChIP-seq data were obtained from Li et al., 2016 (GEO Accession Number GSE81523) (26). Sequencing reads were aligned to a non-repeat-masked version of the *C. elegans* N2 reference genome (ce11) using Bowtie2 with the command bowtie2 —-no-unal —-very-sensitive (27). HSF-1 peaks present at 34°C and their summits were called from Bowtie2-aligned reads with the MACS2 command macs2 callpeak -g ce —-keep-dup auto —-call-summits -q 1e-6 using combined biological replicates and the single input replicate available (28). To normalize ChIP-seq data for display purposes, Bowtie2-mapped reads from combined biological replicates were filtered for duplicates using macs2 filterdup -g ce —-keep-dup auto, and the condition with more mapped reads after filtering was randomly sampled down using macs2 randsample so the total number of reads considered were identical between conditions. Finally, pileup of filtered reads was performed using macs2 pileup with the —-extsize parameter set to the fragment lengths predicted by MACS2 during peak-calling steps.

### HSE identification and scanning

Motifs enriched in 101-bp non-repeat-overlapping HSF-1 summit regions were identified using MEME with the command meme -mod zoops -dna -revcomp (29). The most significant motif identified in HSF-1 peak summits closely resembles the previously-identified HSE motif using the same dataset (26). To scan the *C. elegans* (ce11) genome for HSE locations, MEME-derived output for HSEs was used in conjunction with FIMO and its default parameters, which reports identified HSEs with *p* < 1e-04. HSF-1-bound HSEs were defined as those with at least 14-bp overlap with 201-bp regions centered on HSF-1 summits (30).

### HSF-1 RNAi and overexpression

RNAi was performed by feeding animals either empty vector or *hsf-1(RNAi)*. RNAi experiments were performed as described in (31). The HSF-1 overexpression strain EQ87, *hsf-1p∷hsf-1∷gfp* + *rol-6*, was used for HSF-1 overexpression studies (32, 33).

### qRT-PCR

qPCR was performed as described in (34) except that *ama-1* was used as a reference gene and the Quant Studio machine was used for all experiments. Primer sequences are as follows:

*ama-1*
Forward: CACTGGAGTTGTCCCAATTCTTG
Reverse: TGGAACTCTGGAGTCACACC
*dct-10*
Forward: GTCACACAGCCAACGAATG
Reverse: GTCGGAACTGTACGGATCAT
*Helitron1_CE*
Forward: AATCGTCGTGCCAATACCTC
Reverse: GTGCTCACCGAGATGTCTGA
*hsp-16.2*
Forward: GCTCTGATGGAACGCCAATTTGC
Reverse: CTGTGAGACGTTGAGATTGATGGCAAAC
*hsp-70*
Forward: CCGCTCGTAATCCGGAGAATA CAG
Reverse: CAACCTCAACAACGGGCTTTCC
*linc-7*
Forward: ACCAAGCAGACCCACCCT
Reverse: GTTGATGACGAGACGAGTGTGAG

### Data availability

The RNA-seq datasets generated in this study will be deposited in a public repository upon acceptance of the manuscript. Other data and reagents are available upon request.

## RESULTS

### Multiple classes of RNA genes respond to heat shock in *C. elegans*

The transcriptional response of protein-coding genes (PCGs) to Heat Shock (HS) has been intensively studied (1, 2, 35), yet the effect of this stress on non-coding RNA (ncRNA) expression has not been examined systematically. Here we used RNA sequencing to compare the levels of various types of RNA from last larval stage (L4) *C. elegans* maintained at 20°C (control, CTRL) or shifted to 35°C (heat shock, HS) for 4 (RNA-seq) or 6 hours (smRNA-seq). Along with the expected HS-induced changes in mRNA expression (Supplementary Table S1), this profiling also revealed specific RNA genes belonging to the miRNA, piRNA, lincRNA, ncRNA, pseudogene and repeat families that changed at least two-fold in response to heat stress (Figure 1A).

**Figure 1.**
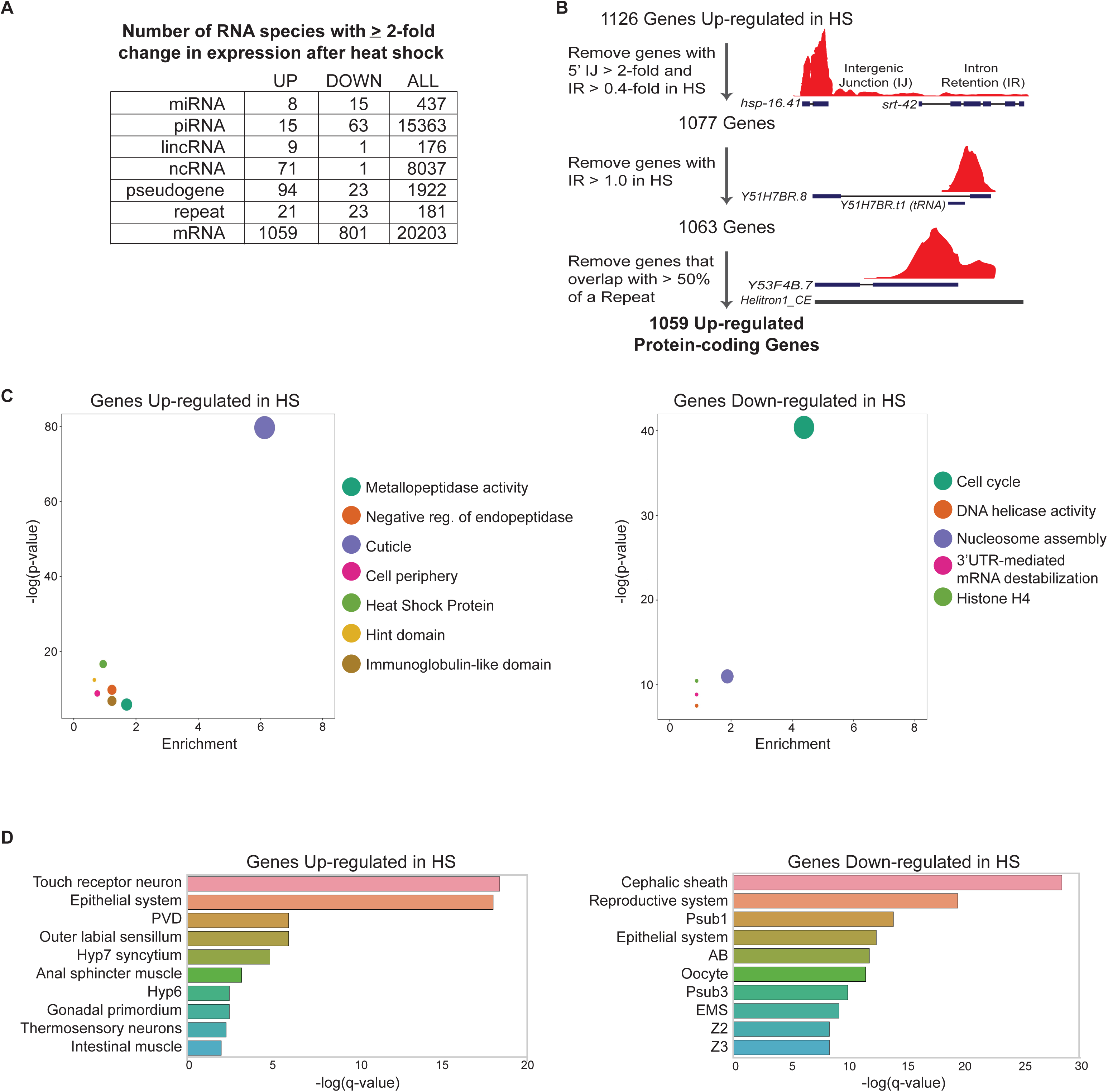
Heat shock alters the expression of coding and non-coding RNAs. (A) Small RNA-seq and stranded paired-end RNA-seq were used to analyze changes in RNA expression in *C. elegans* shifted from 20°C (control, CTRL) to 35°C (heat shock, HS) for 4 (RNA-seq) or 6 hours (smRNA-seq). The numbers of differentially expressed genes in each RNA category are indicated. (B) Strategy used to filter out false positive up-regulated mRNAs. See methods for further details; codes used for filtering are available at https://github.com/wschrein (C) DAVID Functional Annotation Clustering of Genes up- and down-regulated in response to HS (41). Representative members of each cluster with an enrichment score > 2 are shown. Size of dot corresponds to number of genes in each cluster. (D) Tissue Enrichment Analysis (TEA) for genes up- and down-regulated in response to HS. TEA was performed using the Wormbase TEA tool (45). Abbreviations: PVD - Sensory neuron (polymodal nociceptive for mechanosensation and thermosensation), Hyp7 - entire syncytium of hyp7, Hyp6 - Cylindrical hypodermal syncytium in head, Psub1 - Embryonic founder cell, AB - Embryonic founder cell, Psub3 - Embryonic founder cell, EMS - Embryonic Cell, Z2 - Germ line precursor cell, Z3 - Germ line precursor cell.

While our conditions elicited changes in protein-coding mRNA expression comparable to previously published studies of the transcriptional response to HS in *C. elegans* (36, 37), we noticed some peculiarities in the up-regulated gene set. Visual inspection of the sequencing reads mapped to the UCSC Genome Browser indicated several false positive gene calls present in the list of PCGs up-regulated by HS. For example, *srt-42* was originally listed as the second most highly up-regulated gene in HS but reads associated with this gene did not conform to the expected splicing pattern and included sequences stretching upstream into the neighboring heat shock protein gene, *hsp-16.41* (Figure 1B). Thus, the reads covering *srt-42* likely emanate from transcripts that failed to be properly terminated from the highly HS-induced *hsp-16.41* gene. These types of aberrant transcripts, known as DoGs for Downstream of Gene containing transcripts, have been previously detected in various stress conditions (38, 39). While a functional role for DoGs is yet to be demonstrated, incomplete transcriptional termination of a highly induced upstream gene can provide RNA-seq reads that falsely assign a downstream gene on the same strand as also up-regulated (38, 39). Since these false positives are unlikely to maintain coding potential, we filtered them from our original list of HS up-regulated PCGs by removing 49 genes with reads in the upstream Intergenic Junction that were 2-fold higher in HS and that had a ratio of intronic to exonic reads of 0.4 or greater (Figure 1B and Supplementary Table S1).

Another source of mistakenly called up-regulated PCGs was the intronic residence of tRNAs and snoRNAs that apparently failed to be properly terminated in HS (Figure 1B). These longer versions of tRNAs and snoRNAs enabled them to be detected in standard RNA-seq, whereas the normally shorter forms are not captured in this procedure. By filtering out genes with an intron to exon ratio (IR) greater than 1 due to accumulation of an intronic ncRNA in HS, we removed another 14 PCGs from the up-regulated list (Figure 1B and Supplementary Table S1). Finally, PCGs that overlapped a repetitive element that was likely responsible for the increased sequencing reads in HS were also filtered out (Figure 1B and Supplementary Table S1). The bioinformatic steps used for filtering are detailed in the Materials and Methods and available online (https://github.com/wschrein).

With a higher confidence set of PCGs differentially regulated by HS, we confirmed that our conditions elicited changes in mRNA expression comparable to previously published studies of the transcriptional response to HS in *C. elegans* (36, 37, 40). The extensive changes in PCG expression induced by HS aligned with molecular pathways expected to be regulated by this stress. DAVID Gene Ontology (GO) analyses of genes up-regulated by HS indicated strong enrichment for those associated with cuticle maintenance, heat shock proteins, and enzymatic factors (Figure 1C; Supplementary Table S1) (41, 42). Genes in these categories have well-established roles in stress protection and have been previously found to be up-regulated in HS (Figure 1C; Supplementary Table S1) (26, 36, 37). Distinct functional annotations, largely involved in nucleic acid binding, were enriched in the genes down-regulated by heat shock (Figure 1C; Supplementary Table S1). Decreased expression of genes associated with DNA replication is consistent with halted growth and cell division triggered by heat stress (1).

The up- and down-regulated gene-sets also pointed to distinct tissue responses to HS, consistent with cell non-autonomous regulation of the HSR in *C. elegans* (43, 44). Using Tissue Enrichment Analysis (TEA), we observed that up-regulated genes were associated with neuronal and epithelial cells (Figure 1D) (45). In particular, enrichment in thermosensory neurons may reflect the role of these neurons in activating the HSR in a cell non-autonomous manner (Figure 1D) (43). Down-regulated PCGs were associated with reproductive tissues, which likely corresponds to the delay of development in stress conditions.

Having confirmed that the heat shock conditions used in these studies elicited the expected changes in protein-coding gene expression, we next examined the effect of HS on major classes of ncRNA genes. The finding that particular members of each ncRNA class were up- or down-regulated in response to HS suggests that the Heat Shock Response controls the transcription or stability of specific ncRNAs.

### Specific miRNAs respond to thermal stress

Of the 205 miRNAs we detected by small RNA sequencing, 8 were up-regulated at least 2-fold in *C. elegans* subjected to HS compared to CTRL conditions (Figure 2A and B; Supplementary Table S2). Consistent with a previous study, miR-239b was one of the most highly up-regulated miRNAs following HS (Figure 2A-B) (11). The largest fold change was observed for miR-4936, which increased by over 200-fold in HS (Figure 2A-C). This miRNA was virtually undetectable at the CTRL temperature but accumulated to high levels in heat shock (Figure 2C). Furthermore, the predominant isoform we detected for miR-4936 in HS (^5’^ AUUGCUUUGUGGCUUUGCUGGUAAC ^3’^) differs from the reference sequence listed at miRBase (^5’^ UGCUUUGUGGCUUUGCUGGUA ^3’^) (46). This difference in 5’-end sequence is expected to affect target recognition, which is largely driven through initial pairing of miRNA nucleotides 2-8 (seed sequence) (12). Since target mRNA degradation is often the outcome of miRNA regulation, we searched the set of down-regulated PCGs for complementary sites in their 3’UTRs to the seed sequence (nt 2-8) of up-regulated miRNAs (Supplementary Table S2). The predicted targets of these miRNAs reflect GO terms associated with genes whose levels decrease in heat stress, such as 3’UTR binding (Figure 2B). Furthermore, some of these down-regulated genes are potential targets of multiple up-regulated miRNAs, suggesting cooperativity (Figure 2D). These observations are consistent with a role for the up-regulated miRNAs in contributing to the repression of some PCGs in HS.

**Figure 2.**
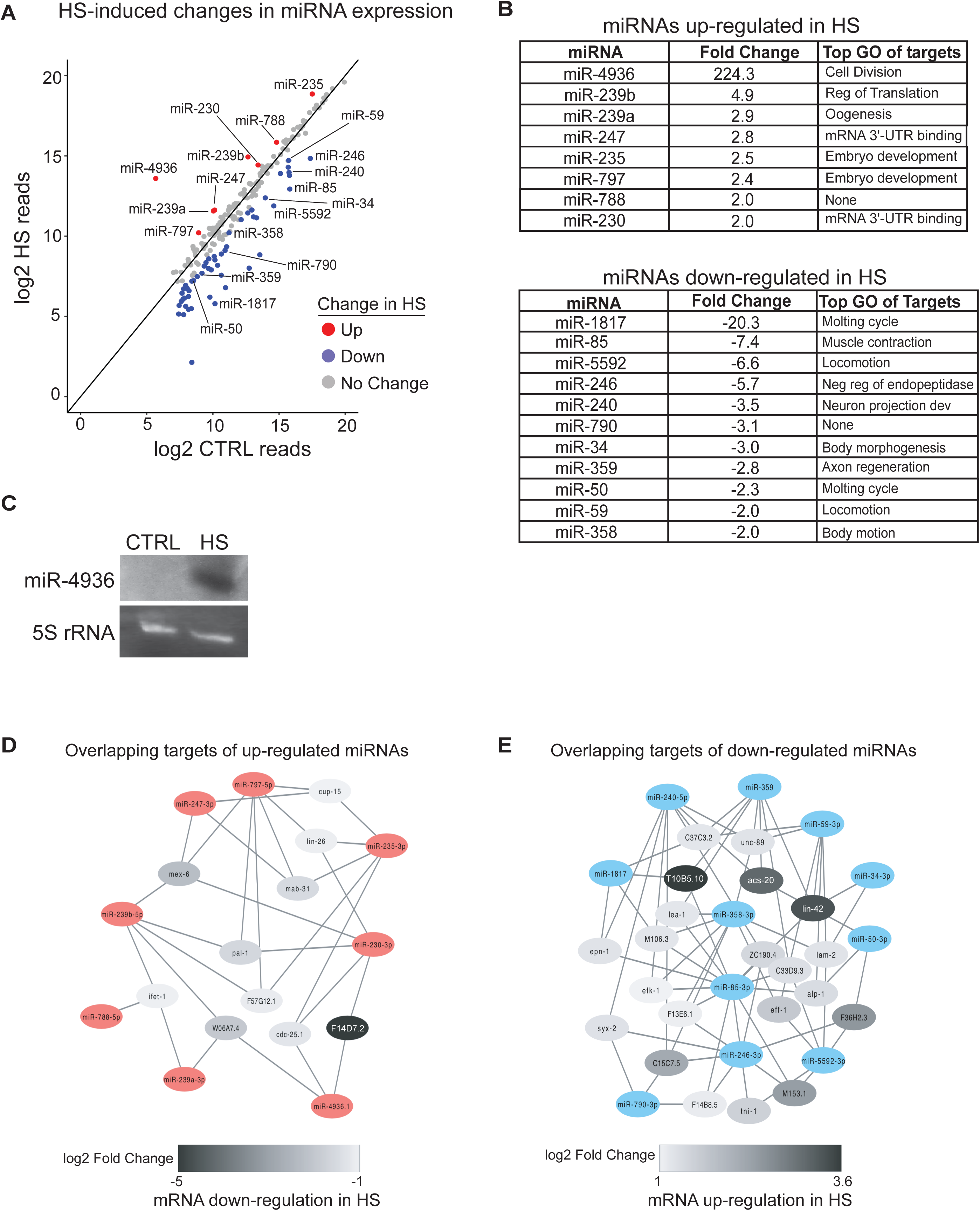
The expression of specific miRNAs is regulated by Heat Shock. (A) Expression of miRNAs in CTRL versus HS detected by small RNA-Sequencing. Results represent the average of two independent biological replicates and miRNAs reproducibly up- (red) and down- (blue) regulated by ≥ 2-fold are indicated. (B) List of guide strand miRNAs within the top 100 expressed miRNAs that are reproducibly up- or down-regulated in response to HS. The final column shows the most highly enriched Molecular Function GO term of potential targets for each miRNA (see Supplementary Table S2). GO analysis was performed using DAVID (41). (C) Analysis of miR-4936 expression in CTRL versus HS conditions by Northern blotting. 5S rRNA serves as a loading control. (D-E) Network analysis of differentially regulated mRNAs targeted by at least 3 up- or down-regulated miRNAs. Cytoscape was used to draw networks (https://cytoscape.org) (25).

Whereas 52 miRNAs were found to be down-regulated in HS, most of these were the lowly expressed passenger strands of the initially processed miRNA duplex (Supplementary Table S2). The levels of only 15 guide strand miRNAs were reduced more than 2-fold after heat shock (Figure 2A-B; Supplementary Table S2). Target predictions for the 11 down-regulated miRNAs that were among the 100 most abundant miRNAs revealed top enriched GO terms associated with molting and movement (Figure 2B). Twenty-one of these target genes have the potential for recognition by 3 or more miRNAs down-regulated by HS, raising the possibility that increased expression of these genes is facilitated by alleviated miRNA repression (Figure 2E). Interestingly, the Period protein homolog *lin-42* is strongly up-regulated in HS and a predicted target of four different miRNAs with reduced expression in HS (Figure 2E; Supplementary Table S2). This gene regulates molting cycles and acts as a general transcriptional repressor of miRNA genes (47–50). Thus, increased expression of *lin-42* in HS could contribute to transcriptional repression of many miRNA genes. This would be consistent with the decreased levels of passenger strand miRNAs, which are more sensitive to changes in transcription than the stable guide strands.

In addition to miRNAs, we also identified 11,979 different piRNAs in our small RNA sequencing data sets. Of these, 15 increased and 63 decreased by at least 2-fold in HS compared to control conditions (Supplementary Table S2). Using the piRTarBase piRNA target prediction tool (51), we found that very few differentially regulated PCGs have potential for regulation by these piRNAs (Supplementary Table S2). Thus, the limited differences in piRNA levels are unlikely to contribute much to the extensive changes in PCG expression observed in HS.

### Specific long non-coding RNAs accumulate in HS

In contrast to miRNAs, where similar numbers of the more abundant species were up- and down-regulated in HS, longer ncRNAs primarily increased in levels in response to heat stress. Approximately 150 genes are annotated as long intergenic non-coding RNAs (lincRNAs) in *C. elegans* (52). While functional roles for the lincRNAs are largely unknown, the lincRNA genes *tts-1* and *rncs-1* have been associated with longevity and stress pathways, respectively (14, 15). Through our RNA profiling, we found that seven relatively abundant lincRNAs increased and only one lincRNA decreased by more than two-fold in HS compared to CTRL conditions (Figure 3A; Supplementary Table S3). Interestingly, *tts-1* was among the highly up-regulated lincRNAs, suggesting that its role in longevity may be linked to a stress pathway (14). At least one of the lincRNAs, *linc-7*, appears to be rapidly up-regulated by HS (Figure 3B). Interestingly, this lincRNA contains five sites with complementarity to the seed region of the rapidly up-regulated miRNA, miR-239b-5p. Additionally, *linc-82* contains eight seed matches for miR-239a-3p and miR-230-3p, both of which accumulate in HS (Figure 2A-B; Supplementary Table S2). These potential lincRNA-miRNA interactions may regulate the stability, availability or function of these specific RNAs during HS.

**Figure 3.**
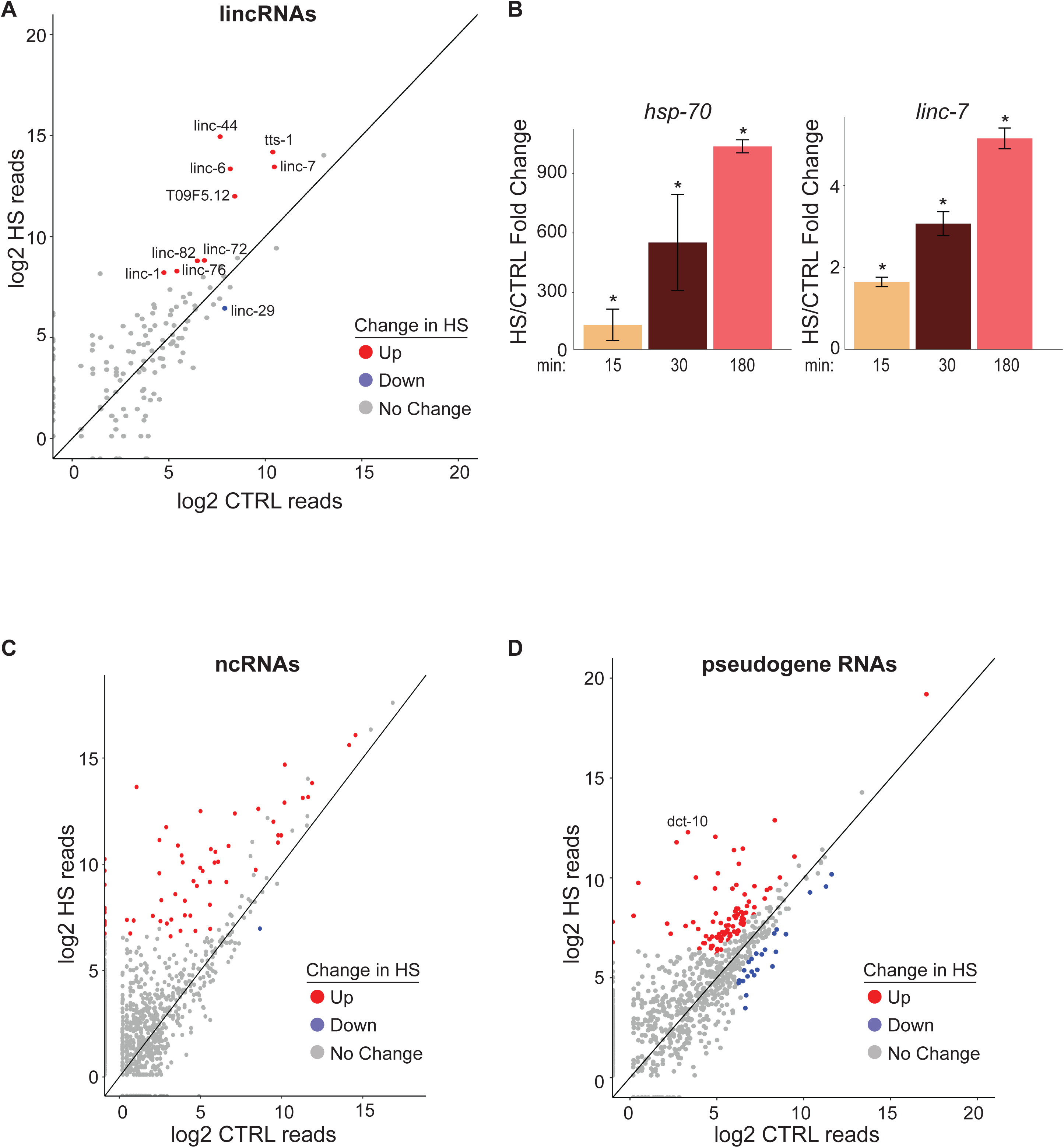
Heat shock alters the expression of long non-coding RNAs (A) Expression of long intergenic non-coding RNAs (lincRNAs) in CTRL versus HS detected by stranded paired-end sequencing. Significantly up- (red) and down-regulated (blue) lincRNAs (≥ 2-fold change with baseMean ≥ 50 and padj < 0.05 from 3 independent replicates) are indicated. (B) qRT-PCR analysis of *hsp-70* and *linc-7* RNA levels after 15, 30 and 180 minutes of HS versus CTRL conditions. Mean fold changes and SEM from 3 independent replicates are shown. \**P* <0.05 (t-test, two-sided). (C) Expression of ncRNAs in CTRL versus HS detected by stranded paired-end sequencing. Significantly up- (red) and down-regulated (blue) ncRNAs (≥ 2-fold change with baseMean ≥ 50 and padj < 0.05 from 3 independent replicates) are indicated. (D) Expression of pseudogene RNAs in CTRL versus HS detected by stranded paired-end sequencing. Significantly up- (red) and down-regulated (blue) pseudogene RNAs (≥ 2-fold change with baseMean ≥ 50 and padj < 0.05 from 3 independent replicates) are indicated.

In addition to lincRNAs, over 4600 ncRNAs that do not belong to a previously characterized class of non-coding RNA are annotated in the *C*. *elegans* genome (22). Of these, 71 increased and 1 decreased at least two-fold in response to HS (Figure 3C; Supplementary Table S3). We also detected expression for 1455 pseudogenes and found that 94 were up- and 23 were down-regulated in HS (Figure 3D; Supplementary Table S4). The relatively large number of pseudogene RNAs that accumulated to high levels in HS suggests that elevated temperatures disrupt the surveillance pathways that normally suppress pseudogene expression.

### Repetitive element-derived RNAs accumulate in heat shock

By re-mapping our RNA-sequencing data to a list of consensus *C. elegans* repetitive elements obtained from repbase (24), we identified reads for 165 types of repetitive elements in the *C. elegans* genome. Similar numbers of repeats (~20) were found to increase or decrease in HS; however, the magnitude of change was greatly amplified in the up-regulated set (Figure 4A-B; Supplementary Table S4). The apparent strong up-regulation of CERP16 is likely due, at least in part, to the position of this element immediately downstream of *hsp-16.2* and *hsp-16.41*. The DoGs produced by each of these highly induced HSP genes are likely a predominant source of reads for this repeat element.

**Figure 4.**
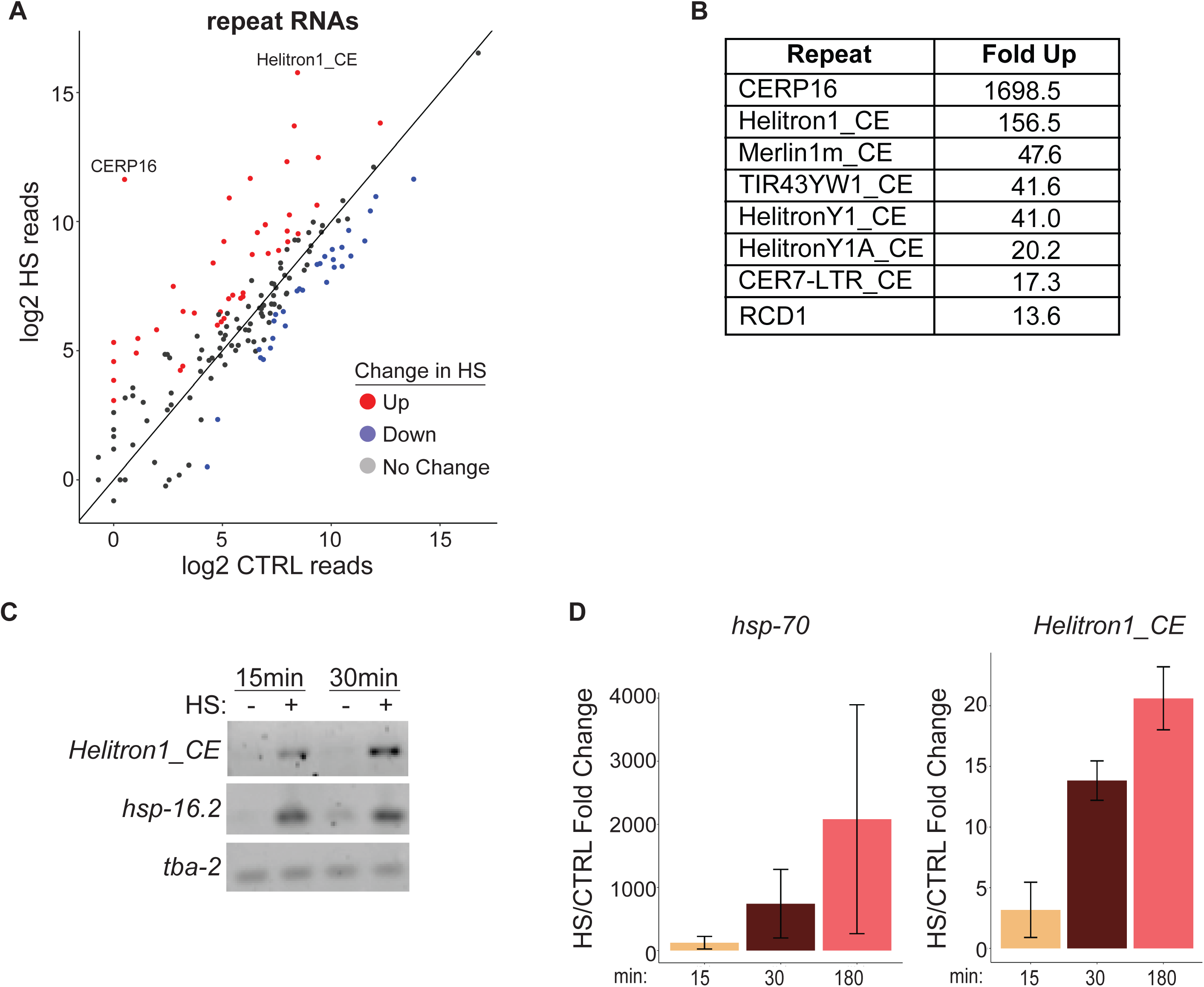
Up-regulation of multiple repetitive element RNAs during heat shock (A) Expression of repetitive element RNAs in CTRL versus HS detected by stranded paired-end sequencing and mapped to a database of repetitive elements (24). Significantly up- (red) and down-regulated (blue) repeat RNAs (≥ 2-fold change with baseMean ≥ 100 and padj < 0.05 from 3 independent replicates) are indicated. (B) List of repetitive element RNAs up-regulated at least 10-fold in HS. (C) Semi-quantitative RT-PCR detection of the indicated RNAs in CTRL and after 15 or 30 minutes of HS. (D) Quantitative RT-PCR analysis of *hsp-70* and *Helitron1_CE* RNA expression during a time course of heat shock. Mean fold changes and SEM from 3 independent replicates are shown. \**P* <0.05, \*\**P* < 0.01, \*\*\**P* < 0.001 (t-test, two-sided).

In contrast, at least some members of the rolling circle DNA transposons known as Helitrons appeared to be strongly induced independently of neighboring PCGs (Figure 4B and see below). RNA from *Helitron1_CE* rapidly accumulated to high levels upon HS (Figure 4C-D). Our reanalysis of previously published *C. elegans* HS RNA-Seq data also confirmed fast induction of Helitron repetitive element RNA (36).

### HSF-1 controls the expression of specific ncRNAs

While defective small RNA silencing pathways may explain some of the increased repeat RNA expression during HS (53), the rapid and massive accumulation of Helitron RNAs suggests that transcriptional induction is also involved. We noted that *Helitron1_CE* exhibited a heat shock response comparable to that of canonical HS-induced genes, such as *hsp-16.42*, with barely detectable RNA levels that quickly rise upon HS (Figure 4C). Thus, we predicted that, like HSP genes, Helitrons could be direct transcriptional targets of HSF-1. To investigate this possibility, we remapped *C. elegans* HSF-1 and Pol II ChIP-Seq data from Li and colleagues to include repeat and other ncRNA genomic loci (26). Similar to the promoter region for *hsp-16.2* and *hsp-16.41*, we found evidence for HSF-1 and Pol II association with *Helitron1_CE*, and other Helitron family members, in response to HS (Figure 5A; Supplementary Tables S1 and S4). Furthermore, the HSF-1 peaks overlapped with multiple copies of Heat Shock Response Elements (HSEs) present in the Helitron genes (Figure 5A; Supplementary Table S4).

**Figure 5.**
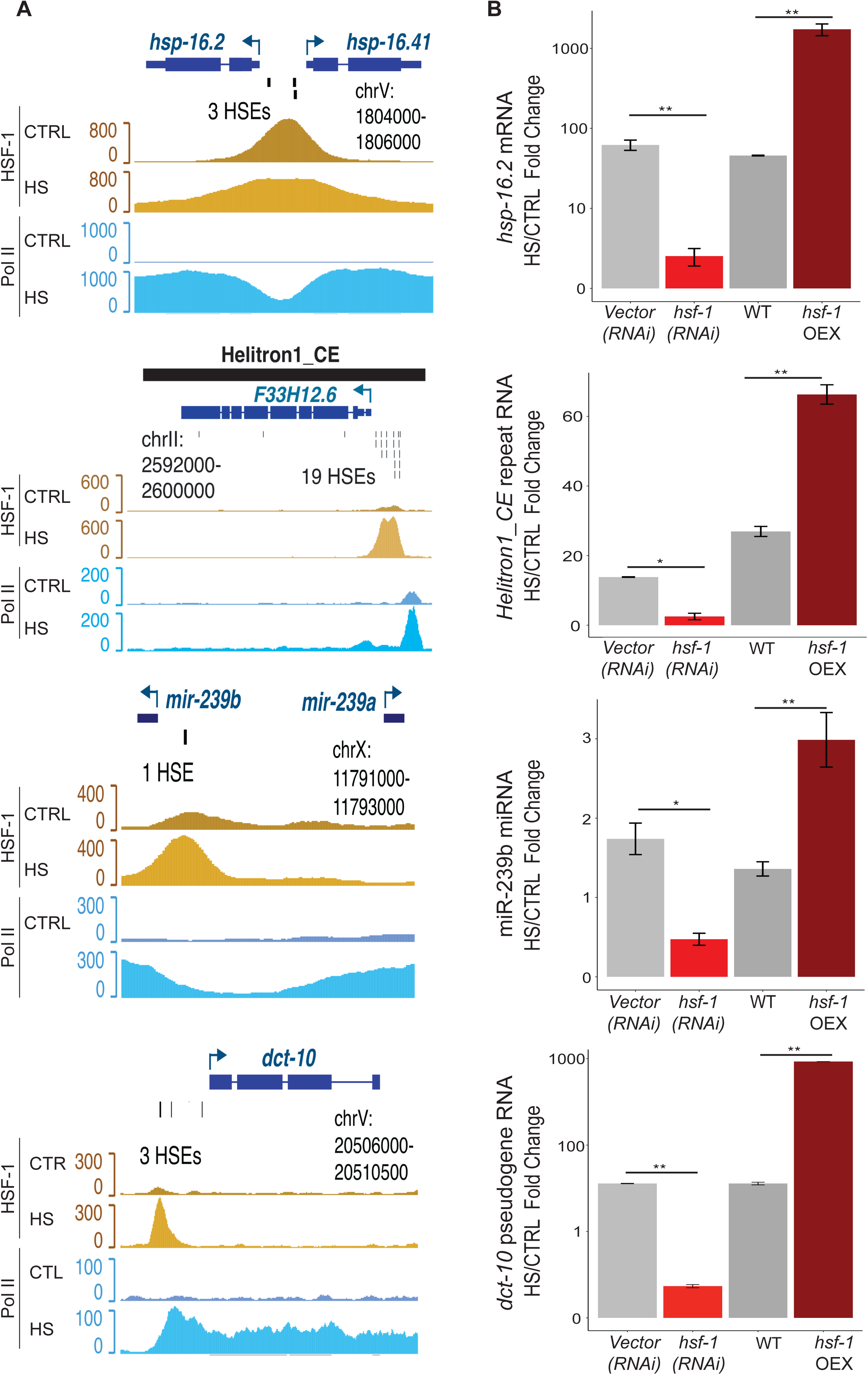
NcRNAs are regulated by HSF-1 during heat shock. (A) Genome browser screenshots of HSF-1 (yellow) and Pol II (blue) ChIP-seq data from control (CTRL) and heat shock (HS) conditions (data from (26)) for representative genes (*hsp-16.2* and *hsp-16.41*, mRNA; *Helitron1_CE*, repeat RNA, *miR-239a* and *miR-239b*, miRNA; *dct-10*, pseudogene). Individual HSEs identified using FIMO (*P* < 1e-04) are indicated (30). (B) Fold change in RNA levels of *hsp16.2, Helitron1_CE, miR-239b*, and *dct-10* after 30 minutes of heat shock in animals subjected to empty vector or *hsf-1* RNAi, and Wildtype (WT) versus a strain overexpressing HSF-1 (*hsf-1* OEX) determined by Quantitative RT-PCR analyses. The mean fold changes and SEM from 3 independent replicates are graphed. \**P* < 0.05, \*\**P* < 0.01 (t-test, two-sided).

The interaction of HSF-1 with Helitron regions suggests that HSF-1 could directly contribute to the up-regulation of Helitron RNA in HS. Using conditions that decrease (*hsf-1(RNAi)*) or increase (transgenic overexpression; *hsf-1*(OEX)) the levels of HSF-1, we observed the expected opposite effects on the expression of *hsp-16.2*, an established HSF-1 target, in response to HS (Figure 5B). Likewise, the induction of *Helitron1_CE* expression by HS was significantly reduced in *hsf-1(RNAi)* and enhanced in *hsf-*1(OEX) conditions (Figure 5B). Given the abundance of HSEs present in some Helitrons and the sensitivity of *Helitron1_CE* to HSF-1 levels, up-regulation of these repeat genes during HS may be largely regulated at the transcriptional level.

To explore the possibility that other ncRNAs may be part of the direct HSF-1 transcriptional program in HS, we analyzed their putative promoter regions (1 kb upstream of the annotated start site) for HSEs and HSF-1 peaks in the ChIP-seq data from Li et al., 2016 (26). Of the up-regulated miRNAs, only the *miR-239* locus fit these criteria (Supplementary Table S2). This region is situated between *miR-239b* and *miR-239a*, which are transcribed in opposite directions. The single HSE and greater level of HSF-1 ChIP-seq reads are closer to the start of *miR-239b*, but enhanced Pol II occupancy in HS is observed over both miRNAs, consistent with their mutual up-regulation in HS (Figures 2A-B and 5A; Supplementary Table S2). Furthermore, we found that reducing or increasing HSF-1 levels resulted in lower or higher miR-239b levels, respectively, in response to HS (Figure 5B). For the up-regulated longer ncRNA genes, HSEs and evidence of HSF-1 binding in HS were detected in the promoter regions of 0 lincRNAs, 11 ncRNAs, and 5 pseudogenes (Supplementary Table S4). Consistent with the DAF-16/FOXO-controlled tumor gene, *dct-10*, being one of the most robustly induced pseudogenes, we found that its promoter contains multiple HSEs and its expression in HS is regulated by *hsf-1* (Figure 5; Supplementary Table S4). Altogether, these findings point to an expanded role for HSF-1 in directing the transcription of specific ncRNA genes as part of the heat shock response in *C. elegans*.

## DISCUSSION

Here we surveyed the response of multiple classes of ncRNA, as well as protein-coding, genes to an episode of HS in *C. elegans*. Our analysis shows that, of the currently annotated genes, HS induced at least a two-fold change in the expression of approximately 9% PCGs, 5% miRNAs, 0.5% piRNAs, 6% lincRNAs, 0.1% ncRNAs, 6% pseudogenes and 24% of the repeat families. Furthermore, some of the most up-regulated ncRNAs, such as miR-4936, were barely detectable under control temperature conditions, demonstrating that often ignored, lowly expressed genes should be reconsidered in different contexts. Our finding that several ncRNA genes parallel canonical HSR genes, such as HSPs, in their dependence on HSF-1 for rapid induction suggests that regulatory RNAs may also have important roles in mitigating the damage caused by excessive heat.

### Aberrant 3’-extended transcripts accumulate in heat shock

During our analysis of differentially regulated PCGs in control versus HS conditions, we noticed a previously described phenomenon known as DoGs for Downstream of Genes (38–40). DoGs result from transcriptional readthrough, leading to mRNAs with long 3’ extensions that include normally intergenic sequence (38, 39). Sometimes DoGs read into adjacent downstream PCGs, which can result in aberrant gene calls (38). Thus, the reads assigned to the downstream gene actually belong to transcripts that are chimeric with the upstream gene and are unlikely to retain coding potential. Such false positives are particularly problematic for compact genomes where closely spaced genes reside in the same orientation. To deal with this issue in our set of PCGs up-regulated in HS, we applied two filters to remove candidates likely emanating from DoGs. We found it necessary to combine the criteria of increased intergenic junction and intron reads because *C. elegans* 5’UTRs are incompletely annotated and splicing is generally less efficient in HS conditions (7, 54) This filtering pipeline may be useful for analyzing other *C. elegans* RNA-seq datasets, since DoGs seem to be generated by a variety of stress conditions (38–40).

The reason for transcriptional readthrough and DoGs generation during stress is currently unclear (38). Since transcriptional termination of most PCGs by RNA Pol II involves the cleavage and polyadenylation machinery, it is conceivable that this process is generally less efficient at elevated temperatures (55). The termination step can also be influenced by chromatin architecture and structural context of the poly(A) signal, which may be sensitive to temperature changes (55). Furthermore, transcriptional termination efficiency by RNA Pol III also seems reduced in HS. We detected 3’-extended tRNA and snoRNA transcripts in the HS, but not control, RNA samples; whereas the mature forms of these Pol III RNAs are too short for capture in standard RNA-Seq library preparations. Since tRNA and snoRNA genes are commonly embedded in introns of PCGs in *C. elegans*, the extended forms observed in HS sometimes overlapped exons. We were able to filter out these falsely called up-regulated PCGs by removing genes with intron retention scores greater than 1. While the cause of 3’-extended Pol II and III transcripts in heat shocked *C. elegans* is yet to be determined, an awareness that these aberrant transcripts can accumulate is important for understanding changes in coding potential induced by stress.

### Specific ncRNAs from multiple classes respond to Heat Shock

Although the majority of individual *C. elegans* miRNA genes are apparently dispensable under laboratory growth conditions, there is accumulating evidence that specific miRNAs play integral roles in a variety of stress response pathways (56–58). Previous studies found that loss of miR-71, miR-246, miR-80, or the miR-229, -64, -66 cluster resulted in increased sensitivity to heat stress (11, 13). We observed that miR-71 is up-regulated 1.4-fold in HS (Supplementary Table S2), which would be consistent with a survival role for this miRNA in elevated temperatures. Conversely, we detected an over 5-fold decrease in miR-246 levels in response to HS, which seems contrary to the profound sensitivity of miR-246 mutants to HS (13). Another counterintuitive change in expression was observed for miR-239a/b. We, and others, detected substantial increases in the levels of these related miRNAs in response to HS (11). Additionally, we present evidence that these miRNAs are transcriptionally induced by HSF-1. Yet, the *miR-239a/b(nDf62)* strain, which lacks both miRNAs, was previously reported to have an enhanced survival phenotype when subjected to HS (11, 13). The identification of direct targets of the differentially regulated miRNAs will be necessary to understand how their altered expression during HS relates to their functional roles.

Most of the miRNAs we found to be up- or down-regulated by HS have not yet been ascribed biological functions. Of the miRNAs that increase in HS, previous reports have documented that miR-247 promotes survival upon exposure to graphene oxide and miR-235 regulates developmental arrest in response to starvation (59, 60). Our observation that these miRNAs increase in HS suggests that they may have roles in multiple stress response pathways. Down-regulation of miR-34 in response to our HS conditions was surprising given its reported roles in promoting dauer survival in response to food deprivation and ensuring normal development in animals subjected to rapid temperature fluctuations (61, 62). However, miR-34 mutants have also been found to exhibit increased and decreased radiosensitivity in the soma and germline, respectively, suggesting that the function of this miRNA is highly context dependent (63).

By far, the most changed miRNA was miR-4936, which increased over 100-fold in HS. Although the dramatic fold change is linked to the virtually undetectable levels in control conditions, this miRNA did accumulate to appreciable levels, as validated by Northern blot. Curiously, this miRNA exhibited more heterogeneity in its mature forms than most other miRNAs. In fact, the predominant species detected in our HS studies differs from the sequence currently present in miRbase (46). While the miR-4936 sequence resides in a predicted hairpin that resembles other miRNA precursors, a corresponding passenger strand has not yet been identified that would be indicative of canonical Dicer processing. Regardless of these peculiarities, it is evident that HS induces massive up-regulation of this RNA, making it a contender for a role in the HSR.

Unlike the comparable numbers of up- and down-regulated miRNAs, members of the longer ncRNA classes predominately increased in HS. Considering that very few functions have been assigned to any *C. elegans* long ncRNAs, the ones that increase in HS are good candidates for potential roles in this stress condition. The lincRNA *tts-1* (transcribed telomeric sequence) has previously been shown to be important for the extended lifespan of animals with reduced insulin signaling (*daf-2* mutants) or mitochondrial activity (*clk-1* mutants) (14). As the levels of *tts-1* are increased in long lived *daf-2* mutants and in animals grown in the presence of bacterial pathogens, the over 10-fold increase in *tts-1* RNA induced by HS could reflect a general response of this lincRNA to stress (14, 64).

Some mammalian long ncRNAs, including pseudogene RNAs, have been assigned roles as competitive endogenous RNAs (ceRNAs) (65). In this role, long ncRNAs have the potential to sequester and sometimes destabilize small RNAs, such as miRNAs, through base-pairing interactions. A caveat to broad regulation of miRNA availability by ceRNAs, is the typically much lower abundance of lincRNAs and pseudogene RNAs (66). Thus, dramatic up-regulation of these types of RNAs, as we observed for specific lincRNAs, ncRNAs and pseudogenes during HS, could boost their regulatory potential. Notably, the up-regulated linc-7 contains 5 sites that support seed pairing to miR-239a-5p and miR-239b-5p. The linc-82 contains repeating elements that can pair with the other half of miR-239a, referred to as the -3p or passenger strand, as well as with miR-230-3p. Considering that these lincRNAs and their potential miRNA partners are all up-regulated in HS, mutual RNA stabilization could be an outcome of these interactions.

There is precedent for repeat RNAs being expressed at higher levels in response to increased temperature (67–70). In *C. elegans*, even a mild temperature elevation can trigger aberrant expression of repetitive loci. Growth at 25°C, instead of the standard condition of 20°C, resulted in increased levels of RNA from some transposons (70). Given the importance of silencing transposons and maintaining genome integrity, it is not surprising that multiple small RNA and chromatin remodeling pathways act to prevent the mobilization of repetitive elements under normal and stress conditions (70, 71). Nonetheless, the up-regulation of RNAs from multiple repeat families during HS indicates that these silencing pathways become compromised in harsh environments. Interestingly, the HS-induced change in repeat RNA expression can be inherited and last for multiple generations in the absence of the original heat stress (70). It has been proposed that certain repeat RNAs or transposon mobilization could be advantageous during stress (72). For example, SINE RNAs in human and mouse cells are induced by HS and act as transcriptional repressors by binding directly to RNA Pol II (73, 74). Additionally, changes in genomic arrangement caused by stress induced expression of transposons could foster beneficial changes in the progeny of stressed parents. Evidence for this idea was recently documented in *Schizosaccharomyces pombe*, where certain forms of stress induced the mobility of transposable elements and the new insertions were linked to enhanced adaptation to the assault (75). As elevated temperatures have been associated with increased genetic mobility in *C. elegans* (76), it is possible that relaxed silencing of transposons during HS could sometimes provide an advantageous genetic change in the progeny of stressed animals.

### HSF-1 regulates the expression of diverse ncRNAs

Activation of the transcription factor HSF-1 in response to HS, and a variety of other environmental perturbations, is essential for driving the expression of factors needed to survive the stress (35). While numerous studies, including our own, have observed the up-regulation of hundreds of PCGs in response to HS (35), recent work in mammalian cells and budding yeast has shown that HSF1/Hsf1 is directly responsible for the transcriptional induction of only a small subset of these genes, which mostly encode molecular chaperones (4, 5). Although it is yet to be established if HSF-1 has a similar limited set of direct essential targets in intact multicellular organisms, it is evident that HS induces widespread changes in HSF-1 binding across the genome in *C. elegans* (26). Here we show that some of these HSF-1-bound loci are associated with ncRNA genes. We identified 2 miRNA, 11 ncRNA, 5 pseudogene, and 5 repeat family genes that were up-regulated in HS and contained HSEs centered within HSF-1 ChIP-seq peaks. Furthermore, we confirmed that the up-regulation of miR-239b, *Helitron1_CE*, and pseudogene *dct-10*, induced by HS was sensitive to HSF-1 levels. A previous study, found that six *C. elegans* miRNAs were dependent on HSF-1 for increased expression in HS (10). Thus, the network of miRNAs, and other ncRNAs, controlled by HSF-1, both directly and indirectly, expands the repertoire of genes regulated by this transcription factor during the HSR. Furthermore, the dramatic, and in some cases rapid, accumulation of specific ncRNAs in response to heat stress in *C. elegans* suggests that functions for some regulatory RNAs may only surface when needed for organismal survival in the natural world.

## ACKNOWLEDGMENTS

We thank Dr. Cindy M. Voisine and members of the Pasquinelli Lab for helpful discussions and critical reading of the manuscript.

